# Neural Mechanisms Associated with Semantic and Basic Self-Oriented Memory Processes Interact to Modulate Self-Esteem

**DOI:** 10.1101/350926

**Authors:** Rachel C. Amey, Jordan B. Leitner, Mengting Liu, Chad E. Forbes

## Abstract

Individuals constantly encounter feedback from others and process this feedback in various ways to maintain positive situational state self-esteem (SSE) in relation to semantic-based or trait self-esteem (TSE). Individuals may utilize a data-driven, or episodic-based process that encodes positive, but not negative, self-relevant information automatically, or employ a semantic-driven process that manipulates encoded negative feedback post-hoc. It’s unclear, however, how these processes work either alone or in concert while individuals receive positive and negative feedback to modulate feedback encoding and subsequent SSE. Utilizing neural regions associated with semantic self-oriented and basic encoding processes (mPFC and PCC respectively), and time-frequency and Granger causality analyses to assess mPFC and PCC interactions, this study examined how encoding of positive and negative self-relevant feedback modulated individuals’ post-task SSE in relation to their TSE while continuous EEG was recorded. Among those with higher levels of TSE, the encoding of positive or negative feedback was not associated with SSE. Rather, higher SSE was associated with mPFC activity to all feedback and higher TSE. The relationship between TSE and SSE was moderated by mPFC-PCC communication such that increases in mPFC-PCC communication led to SSE levels that were consistent with TSE levels. Furthermore, Granger causality analyses indicated that individuals exhibited higher SSE to the extent mPFC influenced PCC in response to positive and negative feedback. Findings highlight the dynamic interplay between semantic self-oriented and basic encoding processes that modulate SSE in relation to TSE, to maintain more positive global selfperceptions in the moment and over time.

## Introduction

Individuals appear inherently motivated to maintain positive self-esteem (SE), an integral aspect of the self-concept [1] and utilize different mechanisms to bolster SE in the face of positive and negative self-relevant feedback [2]. Individuals may initially attend to and encode both positive and negative feedback and then manipulate negative feedback post-hoc in a manner that maintains high SE [3], or they may encode positive feedback more deeply while processing negative feedback at a shallower level, to bias SE in a more positive manner in the aggregate [4]. These self-protection mechanisms suggest a dynamic interplay between more semantic driven self-oriented and basic memory encoding processes. Yet given that many of these findings are based on self-report measures collected long after feedback is received, it’s unclear how and to what extent positive and negative self-relevant information is encoded and processed by individuals on-line in relation to more semantic-based trait SE (TSE) to maintain positive state SE (SSE) in the moment. Employing a social neuroscience approach, the current study examined how individuals encoded positive and negative self-relevant feedback on-line, while they were exposed to said feedback, and how semantic and episodic-based interactions inspired by self protection mechanisms ultimately modulated post-task SSE in relation to TSE. Findings provide evidence for a dynamic interplay between semantic and episodic-based encoding processes that modulates SSE in relation to TSE to maintain more positive global self-perceptions accordingly.

### Social feedback is processed via interactions between semantic self-oriented and basic memory encoding processes

The self-concept is comprised of both abstract information relevant to general self-knowledge (e.g., self-attributes, attitudes and beliefs), or semantic knowledge, and concrete information about past experiences and behaviors, or episodic knowledge [5]. SE, the subjective feelings of one’s self and abilities, has been identified as an integral aspect of the self-concept [1], and likely a product of semantic and episodic self-knowledge. SE theoretically should be affected by positive or negative self-relevant information, which often comes in the form of social feedback. This feedback is essential for evaluating whether situational perceptions are consistent with global perceptions of the self. Prominent theoretical frameworks contradict this assumption. They indicate that individuals have an innate desire to maintain more positive SE in the aggregate [6, 2]. This suggests a potential dynamic interplay between self-based semantic and episodic memory processes. Influential theories pertinent to SE maintenance, including mnemic neglect, positive tropism, and sociometer theory, support this conjecture. Among other things, they argue that situational or SSE may be dependent on self-relevant feedback being processed *post-hoc* in ways that make it consistent with more semantic-based positive global perceptions of the self (or TSE). It has also been argued that SSE may be dependent of feedback received *in the moment* in the form of episodic memory-based encoding biases, depending on the valence of the feedback.

For instance, according to principles of mnemic neglect, given inherent motivations to maintain more positive levels of SE, when individuals encounter positive or negative self-relevant feedback, they utilize basic episodic memory encoding processes to automatically attend to and encode positive information accurately. Negative information, however, is processed at a shallower level [4]. This provides a means for negative self-relevant information to be readily degraded over time and remain distinct from the stored positive self-concept, facilitating positive self-perceptions and SSE in the moment that exist regardless of TSE levels [7].

Conversely, sociometer theory suggests that individuals integrate self-relevant feedback into their self-concept based on situational fluctuations in SSE. When individuals experience initial reductions in SSE they may accurately attend to both positive and negative social feedback via episodic memory-based encoding processes to accurately detect group inclusion and restore SSE [3]. That is, SE serves as a “sociometer” that detects the extent to which one feels included in a valued group. These feelings of exclusion prompt efforts to restore perceptions of inclusion. Sociometer theory also suggests this process may be modulated by TSE. According to Leary, an individual’s “sociometer” may be calibrated differently depending on whether they have high or low TSE. Whereas individuals with low TSE generally feel as though they tend to be rejected, those with higher TSE feel as though they tend to be accepted. As such, individuals with higher TSE may have higher SSE regardless of the type of feedback they are given. This suggests that semantic-based TSE may modulate basic episodic encoding processes involved in on-line conceptualizations of the self, regardless of feedback valence, to maintain SSE in relation to TSE.

Findings from mnemic neglect and sociometer theory allude to a dynamic interplay between semantic and episodic-based encoding processes in response to positive and negative self-relevant feedback during on-line conceptualizations of SSE. Evidence for this potential interplay is informed by an additional theory referred to as positive tropism [8]. According to principles of positive tropism, processing of self-relevant information consists of two phases. Consistent with mnemic neglect, during the first episodic-based phase of evaluation (or the “positive tropism” phase) motivations to maintain SE result in individuals exhibiting an attentional and encoding bias towards positive self-relevant information specifically. During the second stage of information processing, initial information is compared to TSE. Here, positivity biases can be overridden, possibly using more self-oriented semantic processes, if they are inconsistent with TSE [9]. That is, with enough time, semantic-based processes may override basic encoding positivity biases in attention and encoding, suggesting episodic and semantic processes dynamically interact to maintain SSE levels that are consistent with TSE.

Episodic and semantic self-oriented processes thus likely dynamically interact early and often during processing of positive and negative social feedback to influence SSE in relation to TSE. Given that past studies have largely relied on behavioral and self-report measures, it is unclear how exactly these interactions occur. Semantic knowledge about the self may bias how social feedback is encoded or later recalled (and vice versa), but the assessment of self-related measures before or after receiving social feedback ultimately masks how these processes interact. Neuroscience methodologies provide a means to examine this critical time.

### Activity between and within PCC and mPFC can index episodic and semantic interactions in relation to trait and SSE

One way to gain insight into the role that self-based episodic and semantic memory processes play in biasing encoding of feedback and subsequent SSE is to examine activity within and between neural regions that instantiate encoding (i.e., episodic) and global self-oriented (i.e., semantic) processes on-line, while individuals are exposed to positive or negative feedback. Past research has identified a collection of brain regions integral for basic encoding processes including the hippocampus, lateral aspects of temporal cortex, medial and lateral aspects of prefrontal cortex, and the precuneus/posterior cingulate cortex [10]. Of interest to the current study (given its EEG-accessible location in the cortex), the precuneus/posterior cingulate cortex (PCC) has been identified as integral for processes such as attentional allocation, and episodic memory encoding and consolidation [11, 12]. PCC has also been shown to integrate new information in to autobiographical memories during self-reflection [13]. Critically, numerous meta-analyses [10, 14], as well as the fMRI meta-analytic software Neurosynth, highlight an integral role for PCC in basic episodic memory processes among hundreds of studies (e.g., a search for the term “episodic memory” in Neurosynth yielded significant PCC activation, z-score=4.28 among 270 studies) [15].

Conversely, medial prefrontal cortex (mPFC) is considered a hub for semantic based self-oriented processing, including global self-perceptions [16, 17], self-monitoring, the evaluation of one’s direct and reflected self-knowledge [18, 19] and self-esteem. In fact, mPFC has been identified as a potential “neural sociometer” that plays a direct role in SE modulation [20]. mPFC activity has also been correlated with individual differences in SSE and biased perceptions of social feedback estimates, suggesting mPFC may be a hub for the integration of real time feedback with TSE during on-line conceptualizations of SSE [21].

Given the role these two regions play in basic episodic and semantic processing, assessing communication between PCC and mPFC could provide insight in to the degree to which these processes interact during encoding of social feedback and conceptualizations of SSE. Indeed, PCC and mPFC are exclusively recruited (along with other regions in the medial temporal lobe network like the hippocampus) during self-related processes such as autobiographical memory reconstruction and prospection [14, 22], suggesting an intimate relationship between the two with respect to self-maintenance processes in general. Importantly, establishing the temporal directionality of this relationship could provide further insight in to the nature of these interactions. If a given measure of neural activity in mPFC at one time point predicts a given measure of neural activity in PCC at a subsequent time point, this could be indicative of a more semantic-based processing mechanism, whereas the converse could represent a given outcome was driven by more episodic memory encoding processes. Examining these measures of neural function in relation to encoding of information associated with social feedback, SSE, and TSE thus provides a means to clarify the role that episodic and semantic processes play in encoding of social feedback and subsequent SSE.

### Study overview and hypotheses

The current study placed individuals in a context where they received positive and negative social feedback (other supposed students accepting or rejecting them) and then assessed the degree to which individuals encoded information associated with positive or negative feedback (faces of supposed students that accepted or rejected them) while continuous EEG activity was recorded. Individuals then completed SSE measures and a memory test for previously seen faces (as well as lures; unseen faces). Activity within neural regions (measured via power analyses), communication between neural regions (measured via phase locking analyses), and the directionality of this communication (measured via Granger causality analyses) in response to social feedback provided a means to assess the relationship between semantic-based processes, episodic-based feedback encoding, and SSE. Based on mnemic neglect and positive tropism, individuals should exhibit greater memory accuracy for faces associated with positive, as compared to negative feedback. Memory accuracy for positive feedback should be associated with higher SSE. Neurally, greater PCC activity to positive feedback should predict higher memory accuracy for positive feedback and higher levels of SSE in comparison to negative feedback. Based on sociometer theory, individuals may initially encode positive and negative feedback, however, encoding accuracy and PCC activity would not correlate with TSE or SSE. There should only be a relationship between mPFC activity and SE, regardless of feedback encoding and valence.

If these processes interact, more nuanced relationships would be expected. One possibility is that PCC is involved in accurately encoding self-relevant feedback in general (regardless of valence). The effect of feedback encoding on SSE however, is contingent on the degree to which semantic self-oriented processes (i.e., TSE, instantiated by mPFC) become involved to manipulate encoded information to maintain TSE. PCC-mPFC interactions may modulate SSE such that greater mPFC involvement (higher levels of PCC-mPFC communication) is associated with decreased memory accuracy for social feedback and higher levels of SSE. Directionality of this communication could provide further evidence of the influence that semantic (increased connectivity stemming from mPFC to PCC during feedback encoding) and episodic processes (increased connectivity stemming from PCC to mPFC during feedback encoding) have on global conceptualizations of the self in relation to on-line encoding of positive and negative feedback.

## Materials and methods

### Participants

Forty-five white introductory psychology students (23 male, 22 female) participated for partial course credit. All participants were right handed and had no history of concussions, seizures, or brain damage, and had given consent.

### Procedure

The study’s hypotheses were examined using raw data from Leitner et al. (2014) [22]. In that study, participants reported to the lab and were informed that the researchers were ostensibly investigating facial features that promote social interactions. Participants were told that they would be interacting with and receiving feedback from individuals across the country who would view their facial photo on an online social network system. There was no social network and confederate faces were used from the Eberhardt Face Database, Center for Vital Longevity Database [23], MORPH Longitudinal database [24], and from Hehman, Leitner, Deegan and Gaertner (2013) [24]. To corroborate the cover story, participants had their photo taken and were told that it would be uploaded to the social network system. They were then prepared for EEG recording. Next, participants completed the supposed social feedback task (see below) and were led to believe that confederates were deciding in real time whether to accept or reject the participant’s personal profile based on their picture that had just been uploaded to the database. Participants were then given a surprise memory test to measure the extent to which individuals encoded confederate faces that were yoked with either positive or negative social feedback (i.e., accepted or rejected the participant). Finally, participants completed measures of SSE and were debriefed.

### Social feedback task

The social feedback task provided a means to administer adequate amounts of positive and negative social feedback in conjunction with different confederate faces. Each trial began with a fixation cross presented in the middle of the computer screen for 250ms, followed by a confederate face for 2000ms, a black screen for 1000ms, and then feedback indicating whether the confederate accepted them for 2000ms (either “ACCEPT” or “REJECT” was presented in the middle of the screen; Fig 1).

**Fig 1.**
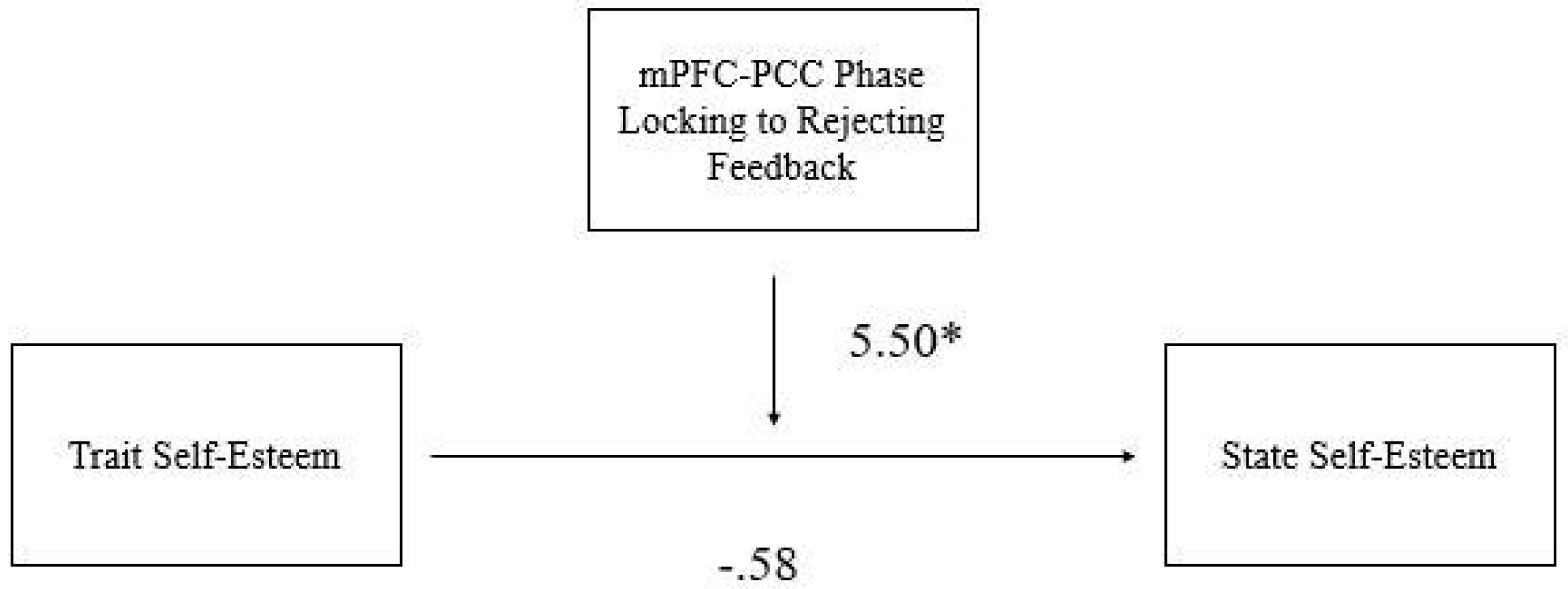
Social feedback task progression.

Participants then were asked to indicate whether the confederate accepted or rejected them with a corresponding button press that would advance them to the next trial. Participants were shown 100 faces randomly paired with feedback (50 faces associated with accept feedback, 50 faces associated with reject feedback). Confederate gender was always matched to participant gender (e.g., males saw only male confederate faces).

### Memory test

Participants were given a surprise memory test consisting of previously seen confederate faces (100 faces; 50 accept, 50 reject) and lures of other confederate faces (50 lures) they had not seen previously. Following the presentation of a confederate face, participants were asked if they had previously seen the confederate on a scale from one to six (where one indicated that they definitely did not see the confederate in the previous task and six indicated that they definitely did see the confederate in the previous task). If participants had been exposed to the confederate during the social feedback task, responses of 4-6 were classified as a hit while responses of 1-3 were classified as a miss. Trials that presented confederates with faces that had not been seen in the previous task were classified as a false alarm if the participant responded with a 4-6, and as a correct rejection if the participant responded with a 1-3. To examine participants’ sensitivity to faces that were associated with social acceptance vs. rejection feedback, separate d’ scores were calculated for faces associated with accept feedback and faces associated with reject feedback. d’, a measure argued to be a more sensitive assessment of memory encoding [25], was derived by subtracting z scores for false alarm rates from z scores for hit rates. Larger d’ values indicate better ability to discriminate seen from unseen faces.

### TSE and SSE

The Rosenberg SE scale (RSE) [26], was administered during a pretesting session conducted at the beginning of the semester as well as post-experiment to compare trait (α=.66) and state (α=.84) SE. The RSE scale given during pretesting to assess TSE framed the questions regarding participants’ overall feelings while the RSE scale given post task to assess SSE framed the questions regarding how participants felt in the moment. Participants answered 10 questions on a 1-4 scale (where one equaled “strongly disagree” and 4 equaled “strongly agree”). Question answers were averaged together. Final scores ranged from 1 to 4 with higher numbers indicating higher TSE and SSE.

### EEG recording

Continuous EEG activity was recorded using an ActiveTwo head cap and the ActiveTwo BioSemi system. Recordings were collected form 64 scalp electrodes. Two electrodes were placed under and on the outer canthus of the right eye to record ocular movements. Data were rereferenced to the original average reference for all analyses off-line and EEG signals were band-pass filtered (.3 to 75 Hz) and stimulus locked to the feedback presentation portion of all trials. EEG signals were stimulus locked to feedback in accordance with previous studies utilizing similar paradigms [23, 29]. All participants’ data were scanned for artifacts using BESA’s artifact scanning tool and ocular artifacts were corrected via the adaptive algorithm implemented in BESA. All participants included in analyses had at least 10 epochs for all trial conditions.

### Source Localization

The goal of this study was not to employ a typical EEG/ERP based approach, which relies on examining evoked activity in response to stimuli within a confined frequency space that collapses across multiple frequency bands and localizing a neural generator for a given ERP of interest. Rather, the current study sought to analyze data in a manner that more closely resembles what fMRI studies typically achieve but with the added benefits afforded to EEG, e.g., examining electrical activity stemming directly from neural activity on the order of milliseconds as opposed to indexing indirect markers of neural activity consisting of blood flow in a given region on the order of seconds. That is, we sought to analyze both spontaneous and evoked activity within specific neural sources across distinct frequency bands thought to reflect different neural processes (e.g., excitatory compared to inhibitory neural processes). To model spontaneous and evoked neural activity, source and time frequency analyses were conducted with Brain Electromagnetic Source Analysis (BESA) 5.3 software (MEGIS Software GmbH, Grafelfing, Germany); MATLAB was utilized for Granger Causality analyses. BESA source localization utilizes a planted dipole approach in which precise coordinates located in specific neural regions are used, as opposed to regions on the scalp, to parse apart the variance in the EEG signal not unlike a typical principal components analytic (PCA) approach (see Scherg et al., 1990 for mathematical proofs) [30]. The primary difference between the two approaches is that whereas PCA is constrained by the mathematical components of the variance within the data, BESA’s approach allows constraints to be based on volume conduction theory and head geometry [30, 31]. This allows one to model principal components as hypothesized sources instead of unique voltage patterns defined by the algorithm. This dipole technique has been cited in notable papers aimed at addressing the inverse problem in EEG source analyses [30, 32] and identifying neural generators of specific ERPs in addition to dipole source localization [33-35].

This a priori hypothesis-driven source localization approach consists of the following steps: 1) electrode space is transformed into a reference free source space. 2) Dipole sources specific to a region of interest are fitted in both orientation and location to best model current flow in a given dipole, independent of other dipoles. 3) This source space is then transformed in to time-frequency space, providing a means to essentially model oscillations in a specific frequency band within a specific source that is theoretically independent of oscillatory activity in other sources. Again, much like the PCA approach, but in this case the principal components are sources with specific coordinates identified in accordance with our hypotheses as opposed to principal components comprised of spatially unique voltage patterns located at the scalp.

Given that prior knowledge of a neural system of interest can be utilized to constrain spatial parameters of a source model [36], our goal was to model and isolate time-frequency activity in sources of interest while also accounting for typical artifacts present in any EEG study (e.g., eye blinks) as well as basic cognitive and perceptual processes that are likely active during any basic cognitive task. Furthermore, because of the social nature of our task, it is possible that other social cognitive processes, e.g., attribution/theory of mind processes, were evoked but irrelevant to study hypotheses as well. Thus, to account for basic visual and perceptual processes like eye movements and eye blinks, multiple sources were initially planted in the left and right eyes, bilateral occipital cortices, and bilateral cerebellum in accordance with previous literature [23, 37]. Dipoles were also planted in regions associated with social interactions and social feedback [38], including the right Temporal Parietal Junction (TPJ, mental state representation, [39, 40]), bilateral Superior Temporal Sulci (STS, social perception, [41, 42]), and Anterior Cingulate Cortex (ACC, social rejection [29, 43]. Final dipoles were then placed in mPFC and PCC in accordance with study hypotheses.

Coordinates for social cognitive processes and hypothesized ROIs were taken from various meta-analyses and relevant works of interest (rTPJ: [44]; right and left STS: [45,46] ACC: [47]; PCC: [13, 48]; mPFC: [20, 44, 49]. These coordinates were then verified using Neurosynth meta analyses to locate the most appropriate talairach coordinates for the model [15]. All sources are plotted in Fig 2 and the corresponding Talaraich coordinates are shown in Table 1. The dipoles from these sources were converted into regional sources with three orientations, each of which was analyzed as a separate dipole. This source model accounted for 98% of the total variance in the EEG signal; no other dipoles could be planted to account for the other 2% in the model. While a model with this many sources in any given brain region would likely account for 98% of the variance, it’s important to note that by applying the approach outlined here, much like a PCA approach and standard fMRI region of interest analysis, we are essentially applying a spatial filter to our data that allows us to examine spontaneous and evoked activity in specific frequency bands at specific points in time in specific ROIs that are theoretically independent of time-frequency activity in other neural regions that are not of interest to the study (i.e., noise). That is, our source model optimized prior knowledge of neural systems to accurately extract signal from theoretically driven ROIs [36].

**Fig 2.**
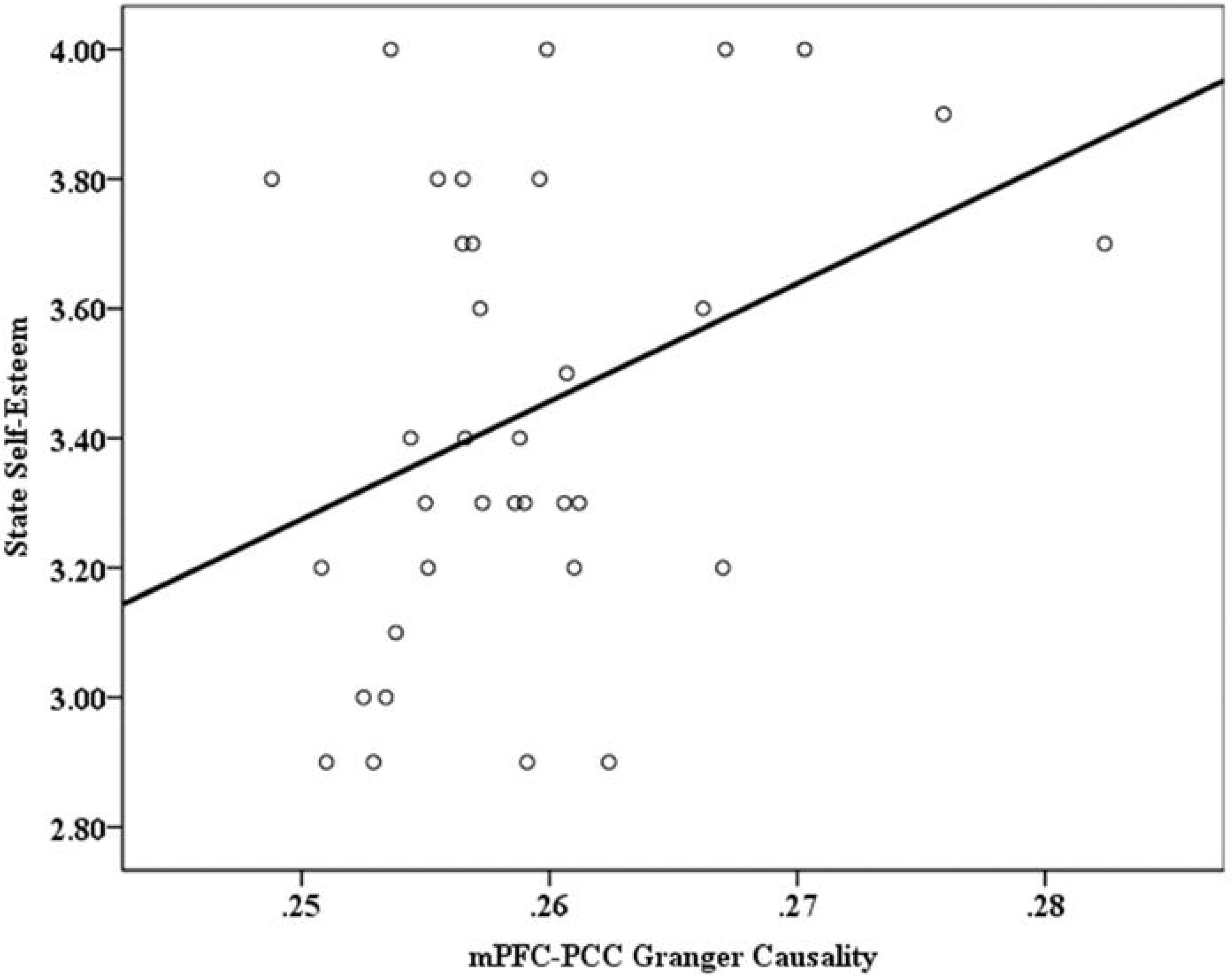
Source model.

**Table 1:**
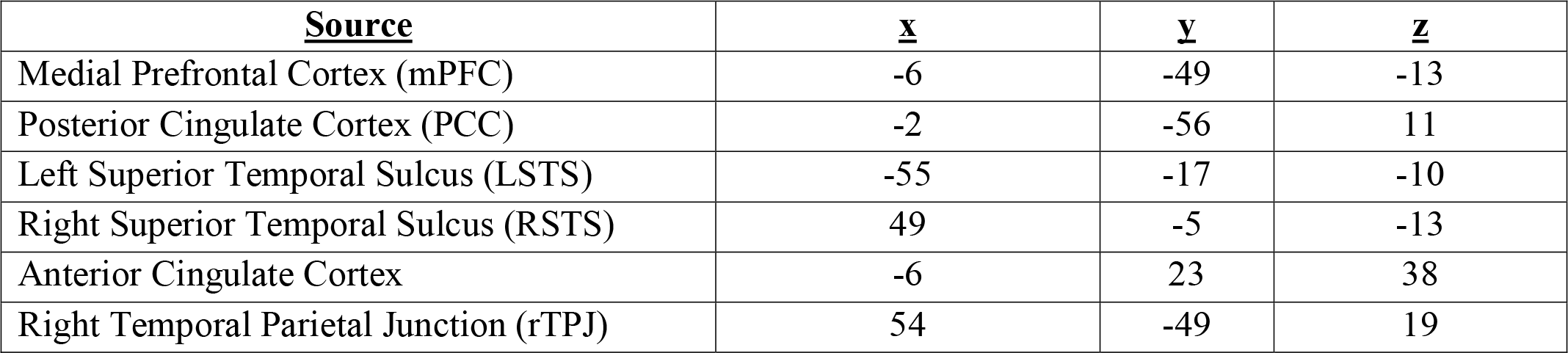
Coordinates for ROIs.

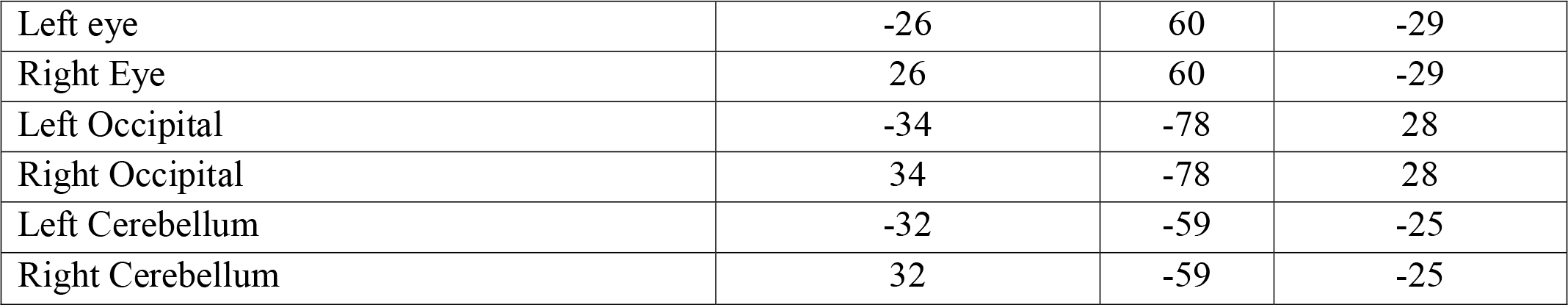

Finally, to validate the source model, a multiple source probe scan (MSPS) was performed in BESA in accordance with past studies using similar techniques [37, 50]. The MSPS model displayed activity around all sources in the source model, suggesting that our source model was an adequate representation of the EEG data.

### Time-frequency analysis

Using the source model above, EEG data were then transformed into time-frequency space using complex demodulation [51] within BESA 5.3. Samples were taken from frequencies ranging from 4 to 50Hz in 2-Hz increments. Theta frequencies were operationalized as 4-8 Hz, alpha as 8-12Hz, beta as 12-30Hz, and gamma as 30-50Hz [52]. Samples were taken from a -500 to 1500ms epoch in 25ms steps. Epochs of interest were extracted from stimulus presentation (0 ms) to 500 ms post-stimulus, specifically from accepting feedback, rejecting feedback, and all feedback presentations (collapsing across accepting and rejecting feedback). This epoch length was chosen for several reasons. One, epochs of 500ms typically helps avoid contaminating the results with edge artifacts. Two, by using a longer time segment it is possible to have better frequency precision and resolution. With a time period of 500ms, two cycles of the lowest frequency of interest (500ms for a 4-Hz oscillation) can be extracted while still capturing higher-frequency activity accurately [53]. Larger time windows are also less susceptible to muscle artifacts, outliers, and other non-brain interference [54].

Phase locking values were obtained for all frequencies above by calculating the correlation of two normalized spectral density functions [55]. All phase-locking analyses used mPFC source as the source reference given its important role in self-related processing. Power was calculated by obtaining the instantaneous envelope amplitude of each source from the model as a function of frequency and latency, following the procedures of past literature [55, 56]. The absolute power in each source with respect to the baseline was then averaged over all trials.

### Time-variant Granger causality

To gain insight in to how self-oriented memory processes interacted, be they basic encoding processes influencing self-oriented processes (PCC to mPFC directionality) or self-oriented processes influencing basic encoding processes (mPFC to PCC directionality), Granger causality (GC) analyses were conducted to assess the directionality of the mPFC and PCC time series. GC is an analytic approach used to quantify the existence and direction of causal influence of time series neural activity from multiple regions. In this study, linear regressive predictive models were first used to calculate independent time series for mPFC and PCC. Time series from either PCC or mPFC were then incorporated into the other using multi-regressive models. If one region has a causal influence on another then the predictive ability of the model should be improved when incorporating the time series from that neural source. See supplemental materials for full mathematical description of the models. In this study, time-variant GC was utilized over a period of 0-500ms post stimulus presentation and more positive GC values represent mPFC activity influencing PCC activity to a greater degree while more negative values indicate PCC activity better predicts mPFC activity.

## Results

### Relationships between trait and SSE

Descriptive statistics for all variables in the study are located in Table 2. An initial linear regression analysis regressing SSE on TSE indicated the two variables were correlated, *b*=.34, t=2.02, R^2^=.11, *p*=.05, Fig 3. Higher TSE (collected at the beginning of the semester) was associated with increases in post-task SSE. It is also important to note that all participants had relatively high levels of TSE (M=3.22, SD=.44, minimum value= 2.4 on a scale of 1-4).

**Fig 3.**
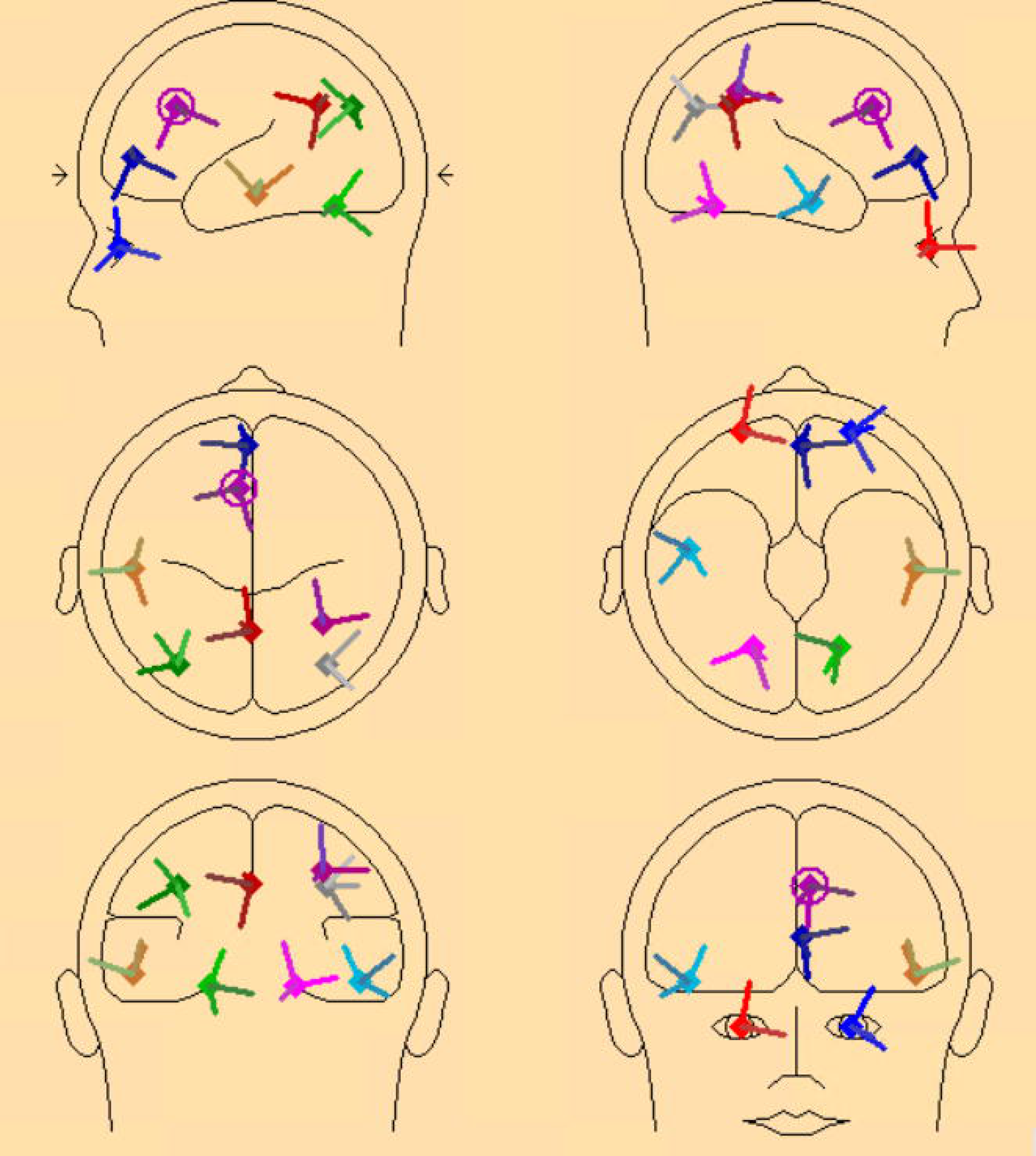
Increases in TSE correlated with increases in post-task SSE.

**Table 2:** Descriptive Statistics of Variables in the Main Analyses.

| <u>Variable</u> | <u>N</u> | <u>Mean</u> | <u>Standard Deviation</u> |
| --- | --- | --- | --- |
| Trait Self-Esteem | 41 | 3.22 | 0.44 |
| State Self-Esteem | 37 | 3.45 | 0.36 |
| Memory Accuracy for Accepting Faces | 44 | 1.56 | 0.65 |
| Memory Accuracy for Rejecting Faces | 44 | 1.61 | 0.65 |
| Overall Memory Accuracy for Faces | 44 | 1.60 | 0.59 |
| PCC Power to all Feedback Theta | 43 | 2005.08 | 1015.46 |
| PCC Power to all Feedback Alpha | 41 | 596.73 | 336.95 |
| PCC Power to all Feedback Beta | 44 | 213.59 | 80.33 |
| PCC Power to all Feedback Gamma | 44 | 109.55 | 66.23 |
| mPFC Power to all Feedback Theta | 41 | 1274.81 | 505.42 |
| mPFC Power to all Feedback Alpha | 45 | 240.79 | 124.97 |
| mPFC Power to all Feedback Beta | 41 | 221.32 | 180.79 |
| mPFC Power to all Feedback Gamma | 40 | 179.60 | 148.43 |
| mPFC-PCC Phase Locking to all Feedback Theta | 45 | 0.15 | 0.05 |
| mPFC-PCC Phase Locking to all Feedback Alpha | 44 | 0.16 | 0.05 |
| mPFC-PCC Phase Locking to all Feedback Beta | 45 | 0.16 | 0.05 |
| mPFC-PCC Phase Locking to all Feedback Gamma | 44 | 0.15 | 0.05 |
| mPFC-PCC Granger Causality | 37 | 0.26 | 0.01 |

### Performance on face memory task

One sample t-tests were conducted comparing participant’s memory accuracy for both accepting and rejecting faces in relation to chance. Both accepting and rejecting faces were significantly different from chance (represented as a d’ score of 0; *p*’s<.001), suggesting that participants reliably encoded accepting and rejecting faces. An initial 2(Gender: male, female) x 2(Face type: accepting, rejecting face) mixed factors ANOVA with repeated measures on the latter variable was then performed on d’ scores (memory accuracy) for accepting and rejecting faces to observe if participants remembered accepting or rejecting faces to a greater extent. This analysis yielded no main effects or interactions (*p*’s>. 09). Thus, participants’ memory for faces did not differ as a function of gender or whether faces were associated with accepting or rejecting feedback, providing little behavioral support for mnemic neglect or positive tropism oriented hypotheses.

### Performance on face memory task in relation to trait and SSE

Regression analyses (collapsing across gender of participant) were conducted to assess whether there was any relationship between SSE and memory accuracy for accepting and rejecting faces independently. Separate analyses regressing state and TSE on to d’ scores for accepting and rejecting faces indicated no relationships or interactions between these variables (*p*’s>.20). These findings provide some initial behavioral support for sociometer theory-oriented hypotheses ([57], study 5) where individuals with higher TSE had higher SSE regardless of whether they accurately encoded who accepted or rejected them during the social feedback task. Encoding confederates who accepted the participant did not predict higher levels of SSE.

Initial behavioral analyses provide some evidence for semantic self-oriented processes biasing basic memory encoding, as the extent to which participants encoded confederates who accepted or rejected them had no effect on SSE respectively, whereas TSE did predict SSE. This suggests that the evaluative nature of the experimental context may have prompted more nuanced interactions between basic encoding and semantic self-oriented processes while individuals encountered social feedback. To gain insight into this possibility, the extent to which mPFC and PCC activity elicited during the feedback task was associated with SE or memory accuracy for faces, and whether interactions between these regions ultimately modulated any of these relationships was explored.

### MPFC and PCC activity and self-esteem

Given that basic d’ analyses indicated individuals did not exhibit a memory bias for accepting or rejecting faces, the remaining analyses collapsed across face type. Analyses specific to accepting and rejecting feedback are located in the supplemental materials. To examine whether MPFC activity was associated with semantic processes, independent linear regression analyses (conducted in all frequency bands for four total models), regressed SSE scores on to mPFC power elicited in response to accept and reject feedback. Because analyses were conducted on four frequency bands, multiple comparisons were controlled for using the Benjamini-Hochberg False Discovery Rate procedure [58], using a q-level of .1 [59-61]. These analyses revealed mPFC power was a predictor of SSE in both beta (*b*=-.48, t=-3.02, R^2^=.23 *p*<.01, SE=.32) and gamma frequency bands (*b*= -.47, t=-2.92, R^2^=.22, *p*<.01, SE=.33). As mPFC power to feedback (both accept and reject) decreased, SSE increased (Fig 4). Theta and alpha frequency bands were not significant (*p*’s>. 11). These relationships were not evident when PCC power (across frequency band) elicited during feedback encoding was entered as the predictor (*p*’s>.24), suggesting these relationships were unique to mPFC power. Identical analyses run in all frequency bands (four independent models), regressed mPFC power elicited during the presentation of feedback stimuli onto TSE. These analyses revealed no relationship between TSE and mPFC power in any frequency band (*p*’s>.20). PCC power also had no relationship with TSE (*p*’s> .21).

**Fig 4.**
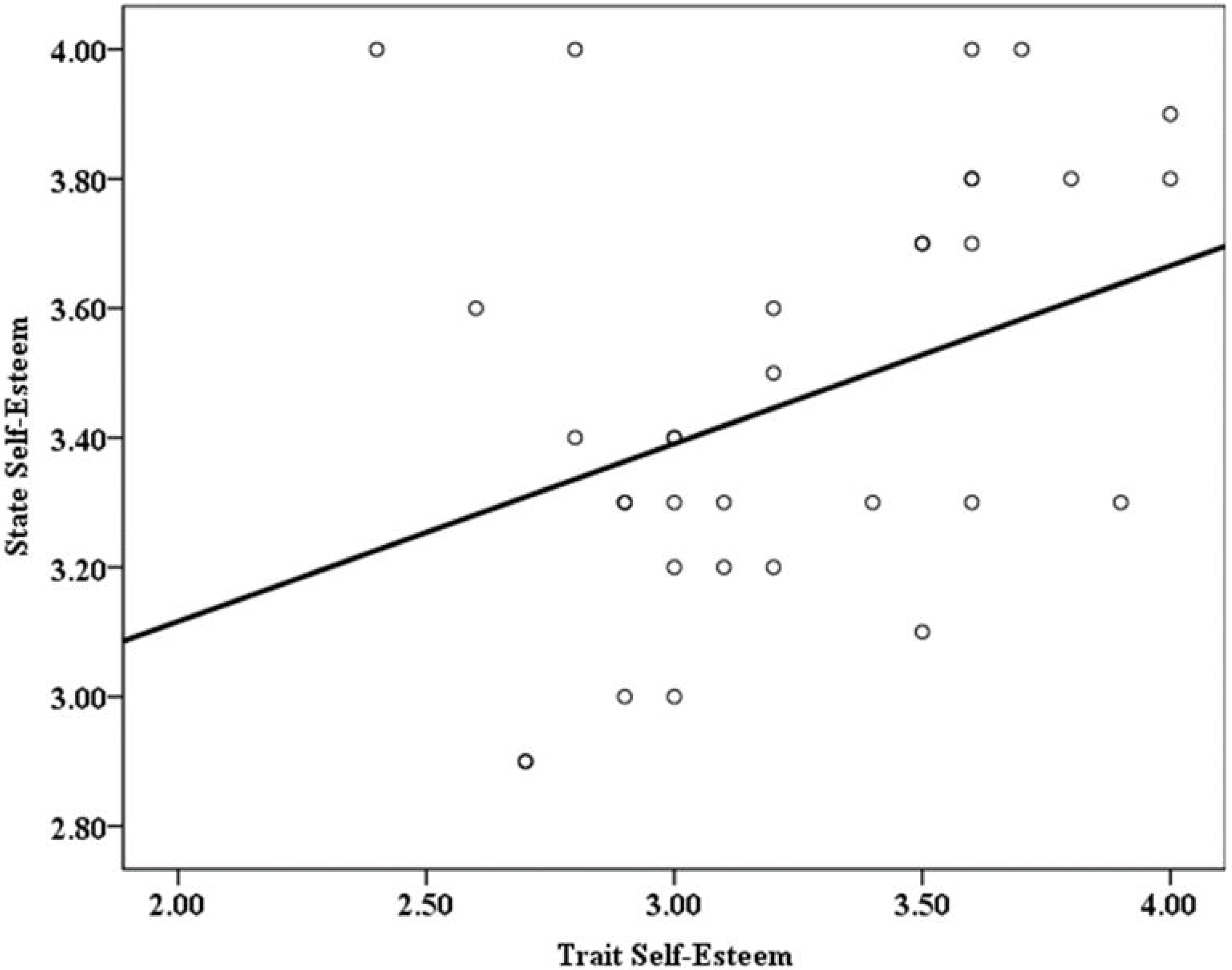
mPFC power in the beta band to self-relevant feedback predicts decreases in posttask state self-esteem.

A final set of analyses regressing mPFC-PCC phase locking values on to trait and SSE were then conducted to examine whether the interaction between these two regions was affected by trait or SSE accordingly. Interestingly, these analyses yielded a positive relationship between mPFC-PCC phase locking in the theta band and TSE, *b*=.36, t=2.38, R^2^=.36 *p*=.02, Fig 5. Participants exhibited more mPFC-PCC phase locking during feedback encoding to the extent they had higher TSE. No other relationships were evident in the other frequency bands (*p*’s>. 12), or between mPFC-PCC phase locking and SSE (*p*’s>.51). These analyses provide some evidence that mPFC was associated with more general SE processes compared to PCC alone and that feedback appeared to play little role with respect to mPFC’s relationship to SE processes (either SSE or TSE).

**Fig 5.**
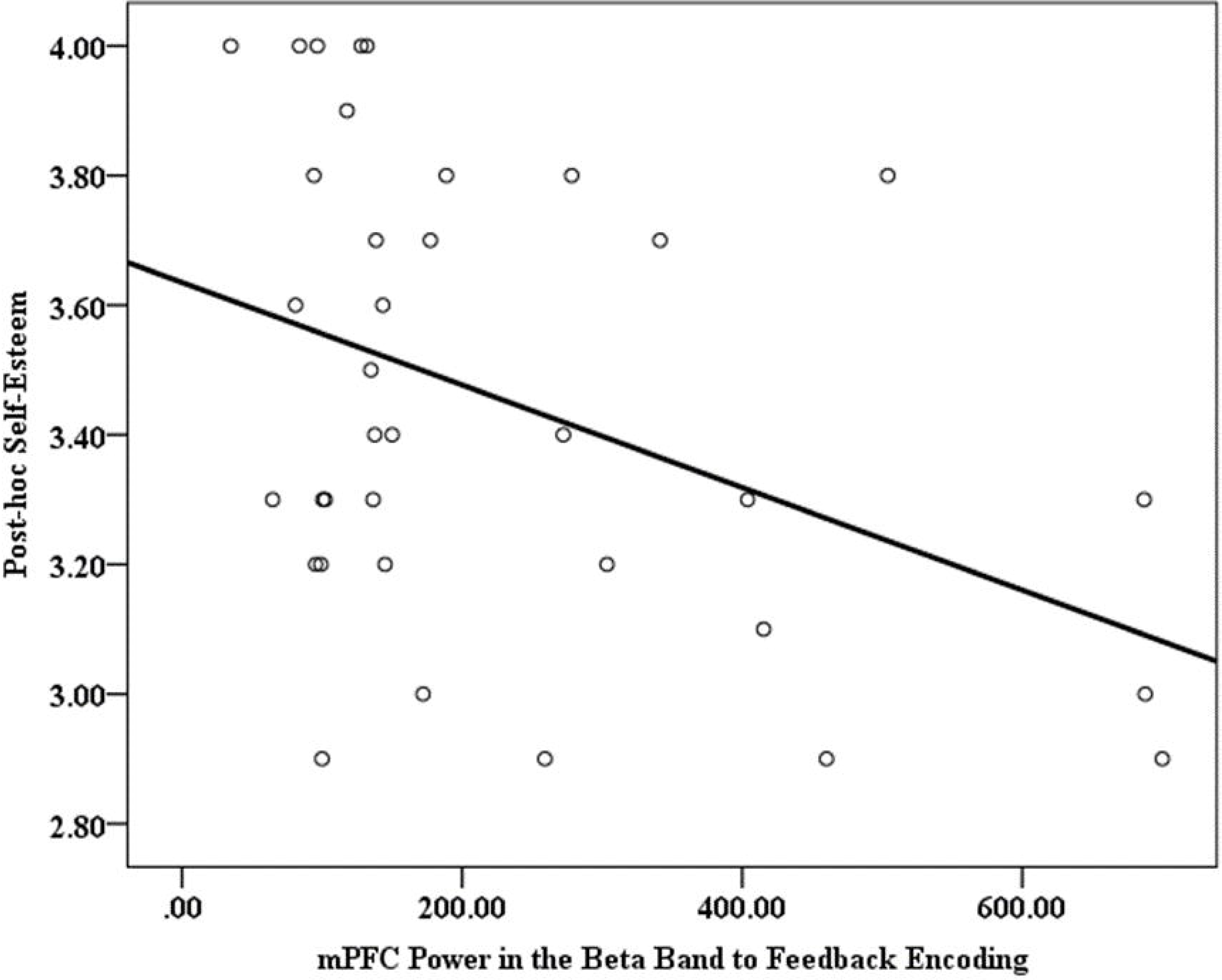
Trait self-esteem positively predicts mPFC-PCC phase locking in the theta frequency band to self-relevant feedback.

### MPFC and PCC activity and memory accuracy for faces

Separate linear regressions (run in all four frequency bands; four models total) were then conducted that regressed memory accuracy for all face types onto PCC power elicited during feedback encoding. Multiple comparisons were controlled for using the Benjamini-Hochberg False Discovery Rate procedure [58], using a q-level of .1 [59-61]. These analyses revealed that PCC power was a significant predictor of accurate memory encoding in the beta frequency band (*b*=.32, t=2.16, R^2^=.10, p<.04, SE=.57). As PCC power to all feedback increased, memory accuracy for all faces increased (Fig 6). After controlling for multiple comparisons this analysis did not reach criterion (p=.034, FDR cutoff p=.025). Nevertheless, given our a priori hypotheses and the large amount of literature supporting this basic finding, we considered these effects to be meaningful. All other frequency bands were not significant (*p*’s>.11). These patterns were not evident when mPFC power (in any frequency band) elicited during feedback encoding was entered as the predictor (*p*’s>.16).

**Fig 6.**
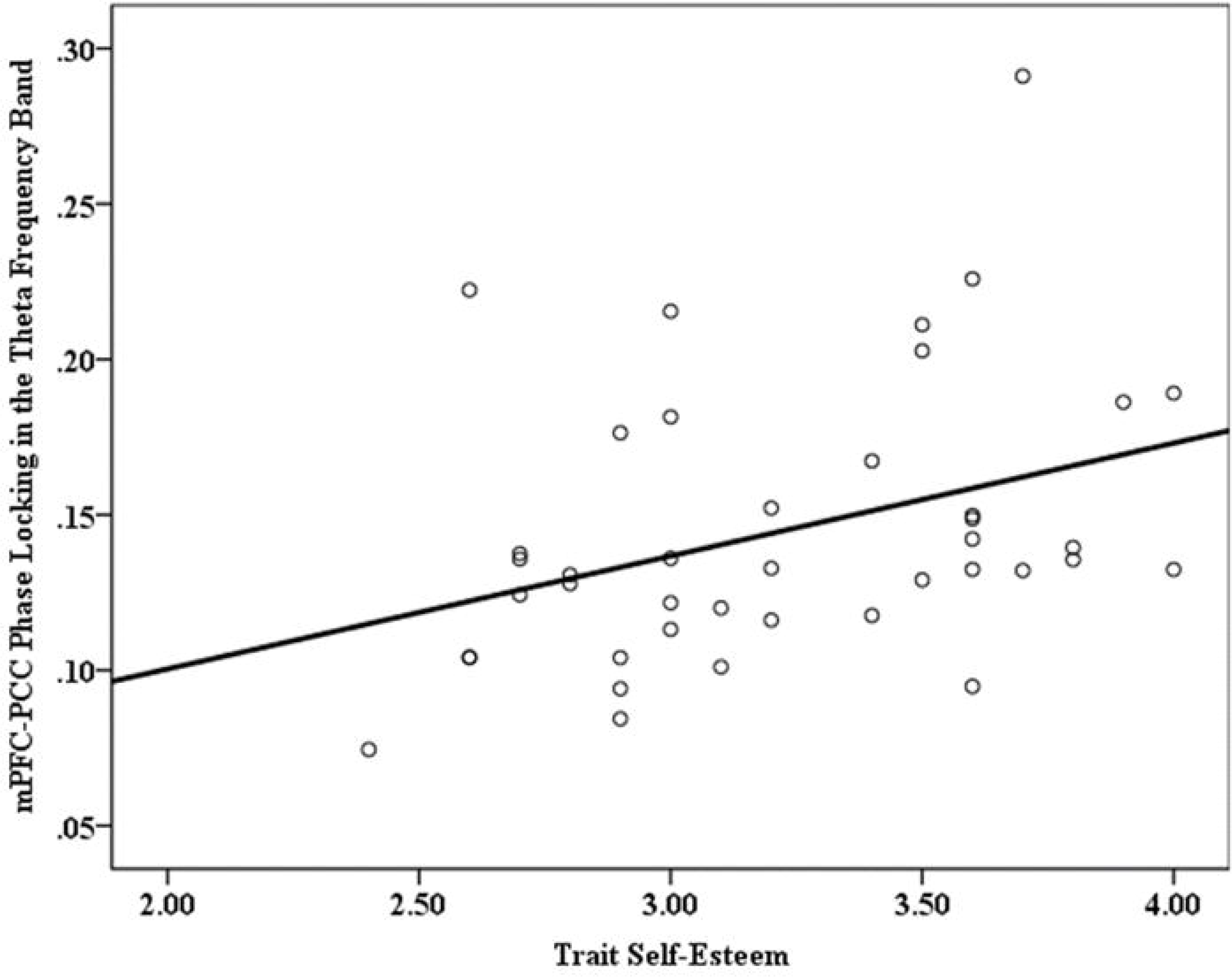
Increases in PCC power in the beta frequency band to self-relevant feedback encoding predicts increases in overall memory accuracy, suggesting an episodic encoding process.

Additional regression analyses regressing memory accuracy for all faces on to mPFC-PCC phase locking elicited in response to all feedback revealed a negative relationship in the beta (*b*=-.45, t=-3.28, R^2^=.20, *p*<. 01, SE=.54) and gamma (*b*=-.30, t=-2.04, R2=.30, *p*<.05, SE=.58) frequency bands. As mPFC-PCC phase locking to all feedback increased, encoding memory accuracy to all faces decreased. Theta and alpha frequency bands demonstrated no relationship (*p*’s>.75). This provides initial evidence that PCC power was associated with basic memory encoding processes, however this relationship was altered when communication between mPFC and PCC were considered.

Thus, whereas PCC power was associated with basic memory encoding processes, and mPFC power was associated with self-oriented processes, the communication between the two regions appeared to be associated with SE and encoding accuracy for all feedback in interesting ways; independent analyses for accepting and rejecting feedback are located in the supporting materials. Individuals exhibited greater mPFC-PCC communication in response to both positive and negative feedback to the extent they had higher TSE, and greater communication between these regions was also associated with decreased encoding accuracy of all feedback (Fig 7). These findings are consistent with past research implicating mPFC and PCC as integral for self-related processing and episodic memory recall and provides evidence that interactions between the two regions underscore a dynamic interplay between semantic self-oriented and episodic processes involved in on-line conceptualizations of the self. Analyses for alternative ROIs are found in the supporting materials, only weak relationships were found with SE and memory providing discriminant validity for mPFC and PCC with respect to SE and memory. The nature of this interaction was addressed next. A discussion of the other sources is located within the supplemental results.

**Fig 7.**
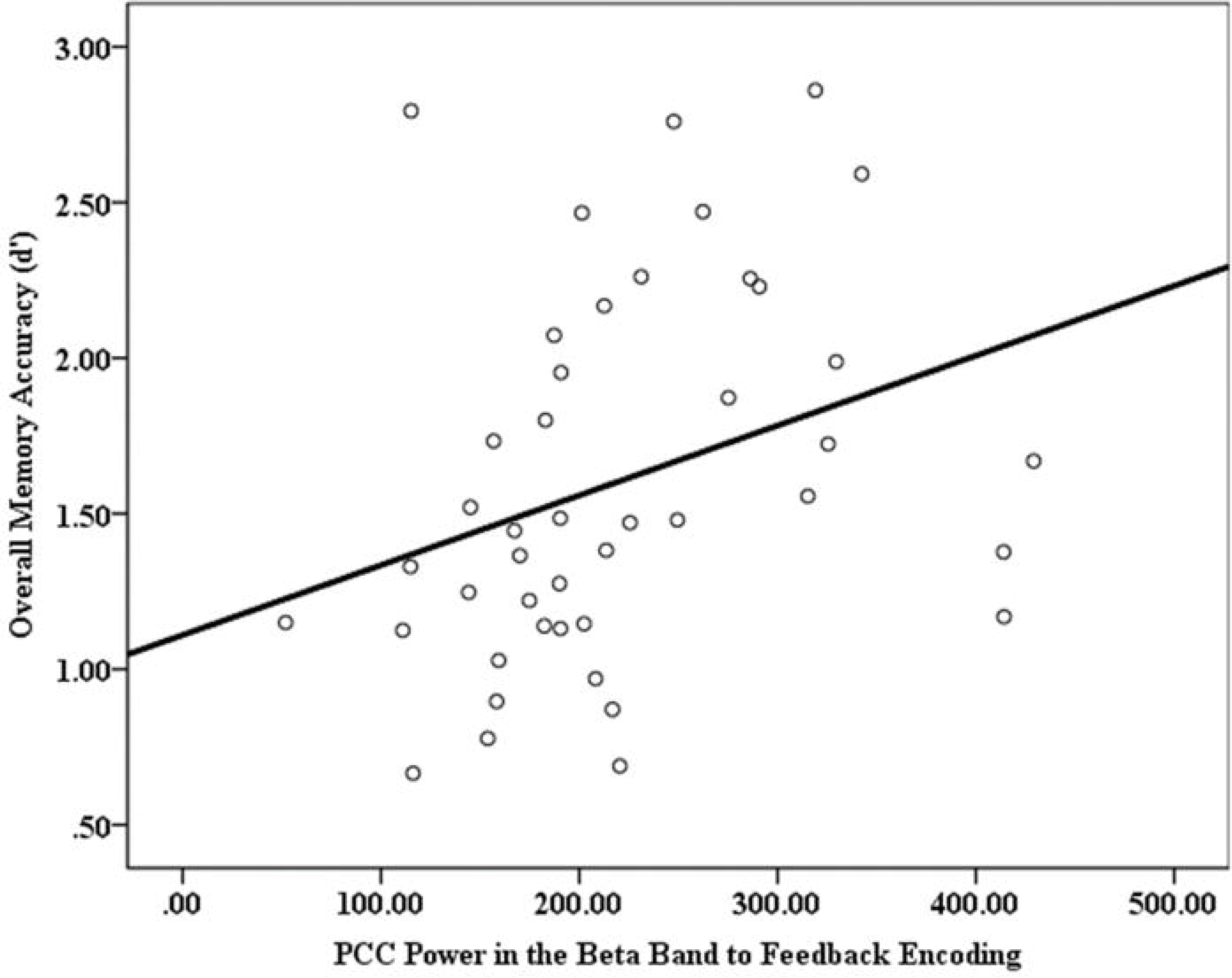
Increases in mPFC-PCC phase locking to self-relevant feedback correlates with decreased overall memory accuracy, suggesting a semantic influence from the mPFC.

### mPFC-PCC communication moderates the relationship between TSE and SSE

Previous literature suggests that TSE may bias encoding of any self-relevant feedback to maintain SSE levels that are consistent with TSE [57]. One way to examine this conjecture is to assess whether on-line communication between mPFC and PCC in response to positive and negative feedback moderated the relationship between TSE and SSE. This possibility was tested via a moderated regression analysis. Moderated regression analyses were tested by deriving unstandardized regression coefficients and 95% bias-corrected confidence intervals (CIs) from 10,000 bootstrap estimates using PROCESS ([62]; model 1). TSE was represented as X, SSE was represented as Y, and mPFC-PCC phase locking to feedback in the gamma frequency band was represented as the moderator, M. Because all four frequency bands were tested, multiple comparisons were controlled for using the Benjamini-Hochberg False Discovery Rate procedure [58], using a q-level of .1 [59-61]. Results yielded a main effect for mPFC-PCC phase locking, *b*=-21.63, t=4.37, SE=1.30, p=.02, such that greater communication between mPFC and PCC was associated with higher SSE scores. mPFC-PCC phase locking also moderated the relationship between TSE and SSE (*b*=6.90, t=2.61, SE=2.64, *p*=.014). Individuals with higher TSE reported higher levels of SSE to the extent that greater mPFC-PCC communication was observed in response to accepting and rejecting faces during the social feedback task (Fig 8). Simple slopes analyses revealed that whereas at lower levels of mPFC-PCC phase locking no relationship was observed between TSE and SSE (*p*=.67), at average and higher levels of phase locking individuals with higher TSE reported higher SSE (*b*_average_=.31, t=2.43, *p*=.0212; *b*_high_=.70, t=3.41, *p*<.01). This pattern was also found in other independent models observing the average and high levels of mPFC-PCC phase locking in beta (*b*_average_=.37, t=2.58, *p*<.02; *b*_high_=.71, t=2.76, *p*<.01) and alpha frequency bands (*b*_average_=.33, t=2.47, *p*=.02; *b*_high_=.60, t=3.08, *p*<.01), and marginally in the theta frequency band (*p*=.11). These results suggest that TSE may influence SSE to the extent that semantic self-knowledge influences basic memory encoding of evaluative social feedback.

**Fig 8.**
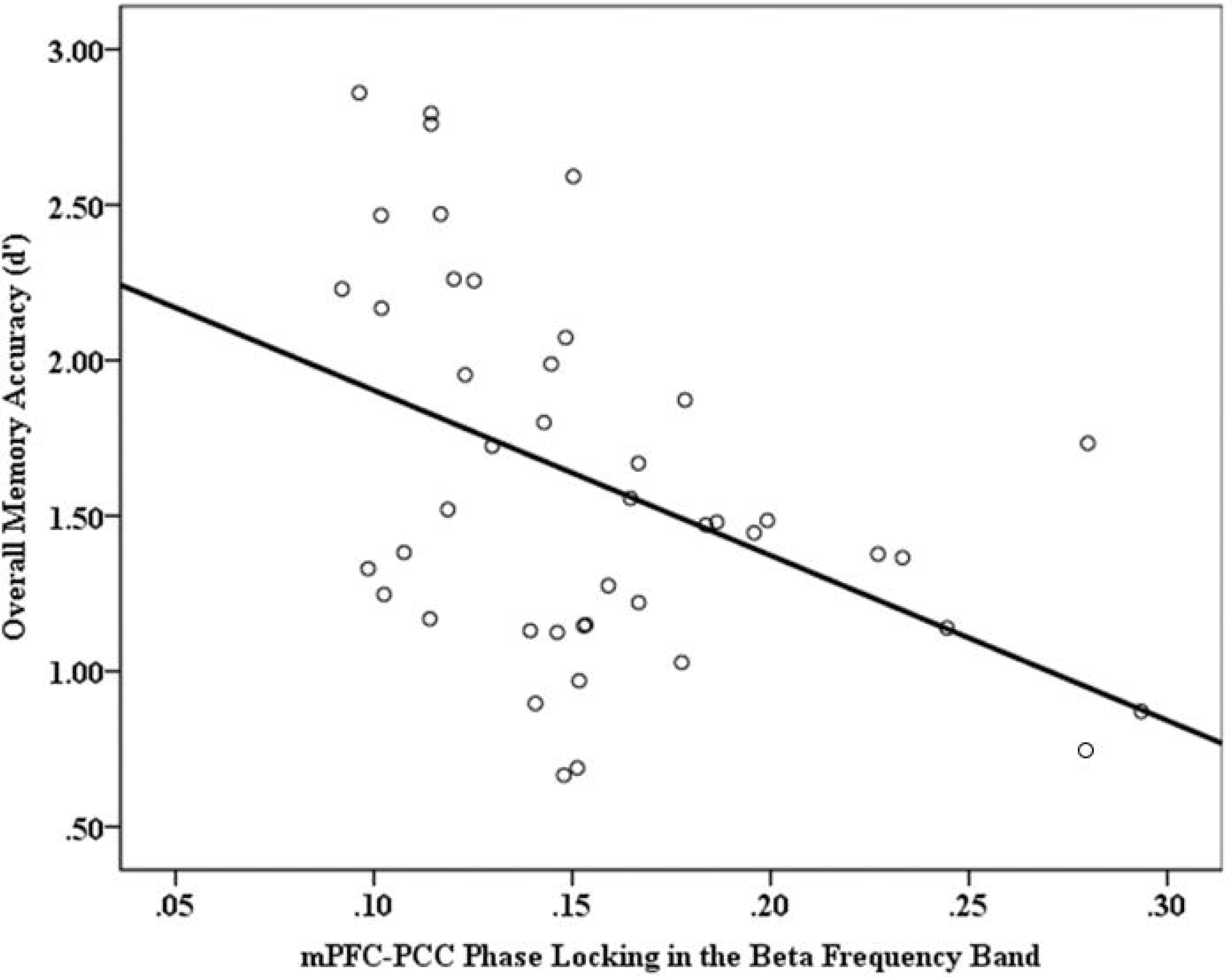
Individuals with higher trait SE reported higher levels of state SE to the extent that greater mPFC-PCC communication was observed in response to accepting and rejecting faces during the social feedback task.

### Exploring the contributions of accepting and rejecting feedback

Although behavioral analyses between accepting and rejecting feedback provided data driven reasoning to collapse across feedback type in the main analyses, according to mnemic neglect, sociometer theory, and positive tropism, accepting and rejecting feedback should be processed differently in the moment to affect the relationship between TSE and SSE. In an exploratory set of analyses, the above moderation was repeated on accepting and rejecting feedback independently to probe for any differences.

### mPFC-PCC communication to accepting feedback moderates trait and SSE

An initial moderated regression analysis (using PROCESS model 1) examined whether on-line communication between mPFC and PCC in response to positive feedback moderated the relationship between TSE and SSE. TSE was represented as X, SSE was represented as Y, and mPFC-PCC phase locking to accepting feedback in the gamma frequency band was represented as the moderator, M. Because all four frequency bands were tested, multiple comparisons were controlled for using the Benjamini-Hochberg procedure [58], using a q-level of .1 [59-61]. Results suggested that the degree of communication between mPFC and PCC towards accepting feedback moderated the relationship between TSE and SSE (p<.05, 95%CI [.1528, 10.8457]), however after correcting for multiple comparisons this effect does not remain.

### mPFC-PCC communication to rejecting feedback moderates trait and SSE

An identical analysis was then conducted utilizing mPFC-PCC communication in response to rejecting feedback (in the gamma frequency band). Results revealed that the degree of communication between mPFC and PCC towards rejecting feedback moderated the relationship between trait and SSE (p<.01, 95% CI [2.1768, 12.6248]; Fig 9) to a greater degree than accepting feedback (R^2^ change_accept_ = .11; R^2^ change_reject_ = .19) and these effects held after correcting for multiple comparisons. Individuals with higher TSE reported higher levels of SSE to the extent that greater mPFC-PCC communication was observed in response to rejecting feedback. Simple slopes analyses revealed that no relationship was observed between TSE and SSE at low levels of phase locking (p=.62). For average and high levels of phase locking, however, TSE positively influenced SSE (*b*_average_=.34, t=2.67, p<.02, 95% CI [.0797, .5960]; *b*_high_=.76, t=3.65, p<.01, 95% CI [.3362, 1.1913]). TSE predicted SSE to the extent mPFC and PCC communicated with one another in response to rejecting feedback. Phase locking in the theta, alpha and beta frequency bands exhibited identical relationships (p_theta_=.09, p_alpha_<.04, p_beta_= .05), however these effects did not survive after multiple comparison corrections. Overall these results provide evidence that exposure to negative feedback drives the relationships between mPFC-PCC communication, TSE and SSE among individuals with higher TSE in general. This suggests self-based semantic processes may play a greater role when high TSE individuals are exposed to negative self-relevant information, possibly reflecting the greater effort that would be required to maintain SSE levels that are consistent with TSE in response to conflicting information. This pattern is consistent with predictions stemming from sociometer theory and positive tropism.

**Fig 9.**
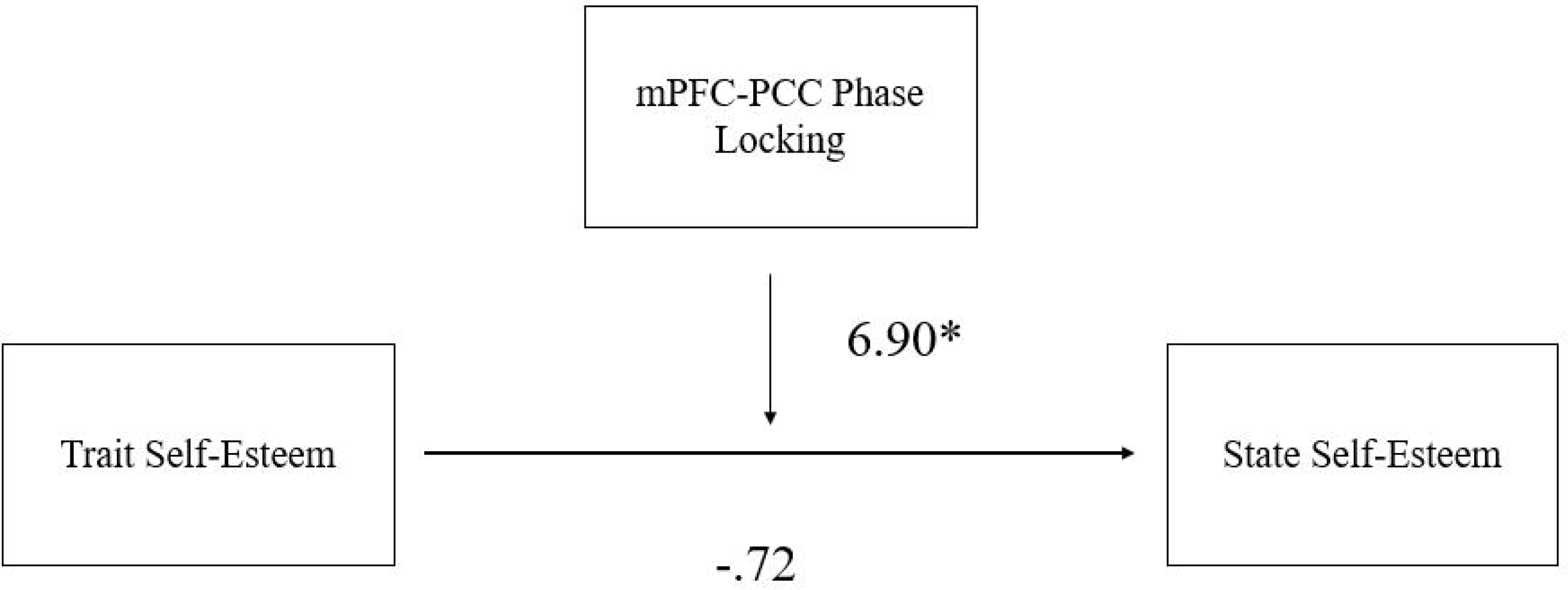
Individuals with higher trait SE reported higher levels of state SE to the extent that greater mPFC-PCC communication was observed in response to rejecting faces specifically during the social feedback task.

### Evidence for the integral role of semantic self-knowledge in self-protective mechanisms

Finally, to determine whether memory accuracy or SSE was driven by more of an influence from semantic self-oriented or basic episodic memory encoding processes, a series of linear regression analyses were conducted utilizing these variables and the product of the time-variant Granger causality (GC) analyses conducted on the mPFC and PCC time series elicited during feedback presentation. Recall that positive GC values represented the degree to which increased activity in mPFC predicted subsequent increases in PCC activity during feedback encoding; higher values indicate a greater influence from mPFC on PCC activity. An initial linear regression analysis regressing SSE onto GC values of mPFC activity predicting PCC activity during feedback encoding revealed that individuals exhibited higher SSE to the extent that mPFC activity predicted PCC activity (*b*=18.19, *p* = .03, R^2^=.13; Fig 10). This relationship was not evident when memory accuracy was modeled as the outcome (*p*>.96). Thus, SSE scores were dependent in part on the extent to which mPFC activity predicted PCC activity in response to evaluative feedback, suggesting semantic self-oriented processes may bias encoding of evaluative feedback on-line to influence SSE in social evaluative contexts.

Fig 10. State self-esteem scores are dependent to the extent to which mPFC activity predicted PCC activity in response to self-relevant feedback, suggesting a semantic encoding mechanism.

## Discussion

By examining how brain and behavioral measures associated with semantic-based self-knowledge and episodic encoding processes interact in response to social feedback, findings from the current study inform our understanding of how these processes interact to affect SSE in relation to global self-perceptions and TSE. Consistent with past research identifying mPFC as a hub for self-oriented processes, like SSE and TSE ([16,20]), mPFC activity (power) elicited in response to positive and negative social feedback predicted SSE but not memory accuracy for faces associated with social feedback. Conversely, consistent with previous research citing PCC as an integral component in basic memory encoding [11] PCC activity (power) in response to positive and negative social feedback was associated with increases in memory accuracy for both accepting and rejecting faces, but not trait or SSE (although this effect was slightly less reliable considering multiple comparison corrections). These relationships were not evident in other sources identified in source localization analyses (e.g., TPJ, STS, ACC), providing evidence for discriminant validity among mPFC and PCC with respect to self-based semantic and episodic encoding processes accordingly.

Importantly, findings from the current study allude to a dynamic interaction between self-based semantic and episodic memory processes with respect to chronic self-perceptions (TSE) and “actual data”, i.e., self-relevant information received in an evaluative context. Moderated regression analyses revealed that individuals with higher TSE exhibited more mPFC-PCC phase locking while being exposed to social feedback. Greater communication between these regions, in turn, predicted a positive relationship between TSE and SSE, which was driven mPFC-PCC communication in response to negative feedback specifically. Highlighting how self-oriented processes can directly influence basic encoding to affect SSE, GC analyses indicated that SSE was higher to the extent mPFC power had a direct, causal influence on PCC power during individuals’ exposure to feedback.

Consistent with Leary [57], the parameters under which mPFC becomes involved in PCC-based encoding may be influenced by the context and valence of TSE and SSE. According to Leary, individuals with higher chronic TSE maintain higher levels of SSE by assuming they will be accepted in a given context and thus do not attend to any kind of self-relevant feedback in general. This argument is consistent with findings from the current study indicating that when TSE was higher, encoding processes did not exhibit any relationships with SSE; individuals’ SSE was consistent with their TSE regardless of whether they were socially rejected or accepted.

Although relationships between mPFC-PCC communication to accepting feedback, TSE, and SSE were evident, it was the communication between these regions in response to rejecting feedback that appeared to play a larger role in TSE-based maintenance of SSE. These findings support hypotheses derived from sociometer theory and positive tropism, which suggest that negative feedback may be manipulated post-hoc among high TSE individuals to maintain high SSE accordingly. Results extend the understanding of these mechanisms by suggesting that the influence and nature of basic encoding is predicated on individuals’ semantic-based TSE while individuals receive self-oriented feedback. They also provide evidence that actual data is in fact encoded, at least with respect to the PCC, but the dynamic interplay between mPFC and PCC may ultimately bias how this data is factored in to global conceptualizations of SSE, which is something all three theories have not clearly explained or expanded upon to date. Importantly, findings from this study help bridge the gap between past research that primarily examines SSE via pre and post self-report measures independent of what individuals encode in a given context, and social neuroscience studies that did not examine how multiple regions integral for semantic self-knowledge and episodic encoding processes interact and/or directly influence one another to affect SSE with respect to TSE.

While no a priori hypotheses were established for specific frequency bands, it’s worthwhile to note that different findings appeared to be driven by activity in beta and gamma frequency bands. Whereas analyses focusing on activity within regions typically found effects in the beta and gamma frequency bands, some phase-locking analyses suggested effects were specific to the theta frequency band (although most analyses provided evidence that all frequency bands were trending in the same direction except for the alpha frequency band). Clearly there is still much debate in the literature regarding what exactly different frequency bands correspond to neurally and psychologically, however, findings from this study are rather consistent with current theoretical accounts. Neuronal oscillations are an essential part of the brain’s design, suggesting functional relevance for each frequency band [63]. Frequency bands have been associated with different brain states and processing mechanisms including attention and memory [64, 65], and can be characterized by the neural area activated. For instance, higher frequency oscillations have been considered representative of smaller neural networks in a given cycle (i.e., activity within a brain region) while lower frequency oscillations have been considered representative of larger network interactions (i.e., activity between brain regions; [66]). Ultimately all frequency bands have been shown to temporally coexist within the same neural structures, however, as a given neural structure performs operations locally as well as in relation to larger networks. These patterns of oscillations and functional architecture of the brain allow multiple cognitive processes to be carried out at once, possibly in a hierarchical manner [64].

Thus, with respect to the findings, analyses confined to activity within a region or driven by a single region (e.g., like those found for mPFC and SE and PCC and memory accuracy) would be expected to yield meaningful results in higher frequency bands such as beta or gamma. Likewise, analyses focusing on activity between regions (e.g., mPFC-PCC phase locking analyses) should yield meaningful results in lower frequency bands like theta, which is consistent with findings from this study.

Specific to study limitations, as with any EEG study that utilizes a source localization approach, it is always important to encourage caution with respect to conjectures based on specific regions of the brain given limitations in spatial localization associated with the methodology. Nevertheless, standards practiced in this study, e.g., using a high density EEG array for data collection and restricting sources to outer cortex, have been shown to provide fairly precise measurements of specific brain regions (with EEG regional source voxel clusters around 7 cubic millimeters in size as opposed to 3 cubic millimeters for fMRI; [53]). Given the number of sources present in our model it is also possible that any combination of sources could provide a good representation of the global EEG signal. While true, a theoretical approach was taken in selecting these sources. Prior knowledge of regions integral in social cognition were used to constrain spatial parameters of the model [37]. It is also important to note that only a priori defined mPFC and PCC sources exhibited any hypothesized relationships with behavioral outcomes of interest. No less, these behavioral outcomes map directly onto hypothesized functions of the a priori regions of interest. That is, mPFC activity predicted self-esteem but not memory measures and PCC activity predicted basic memory but not self-esteem measures, providing an element of construct and discriminant validity to our basic findings. MSPS analyses bolstered confidence in source locations by revealing activation around a priori regions as well. Nevertheless, future research should replicate this study utilizing combined EEG-fMRI methodologies to allow for both optimal temporal and spatial resolution to bolster claims accordingly.

In summary, findings from this study highlight the parameters under which self-oriented semantic and episodic memory processes interact to modulate SSE in relation to TSE. When TSE is high, mPFC appears to play an integral role in altering how social feedback is recalled, such that memory accuracy for socially accepting and rejecting individuals does not appear to influence the extent to which higher SSE is consistent with higher TSE. This suggests semantic-based TSE modulates the extent to which basic encoding processes are employed in evaluative contexts, with the goal of ultimately maintaining SSE levels that are consistent with TSE levels. Thus, findings provide additional support for theories of self-maintenance that argue for the importance of semantic and episodic processes, but extend upon these theories by suggesting the self-maintenance process is much more dynamic and nuanced than typical self-report measures would suggest.

## Acknowledgements

The authors would like to thank all Forbes Lab research assistants for helping with data collection and organization.

## Supporting Information

**S1 File. Supporting analyses and results.**

**S1 Table. Descriptive statistics for supporting analyses.**

**S2 Table. Descriptive statistics for alternative ROIs.**

## References

1. Greenwald AG, Bellezza FS, Banaji MR. Is self-esteem a central ingredient of the self-concept?. Personality and Social Psychology Bulletin. 1988 Mar;14(1):34–45.

2. Tesser A. On the confluence of self-esteem maintenance mechanisms. Personality and Social Psychology Review. 2000 Nov;4(4):290–9.

3. Leary MR, Baumeister RF. The nature and function of self-esteem: Sociometer theory. In Advances in experimental social psychology 2000 Jan 1 (Vol. 32, pp. 1–62). Academic Press.

4. Sedikides C, Green JD. The mnemic neglect model: Experimental demonstrations of inhibitory repression in normal adults. Behavioral and Brain Sciences. 2006 Oct;29(5):532–3.

5. Kihlstrom JF, Albright JS, Klein SB, Cantor N, Chew BR, Niedenthal PM. Information processing and the study of the self. In Advances in experimental social psychology 1988 Jan 1 (Vol. 21, pp. 145–178). Academic Press.

6. Leary MR, Haupt AL, Strausser KS, Chokel JT. Calibrating the sociometer: The relationship between interpersonal appraisals and the state self-esteem. Journal of personality and social psychology. 1998 May;74(5):1290.

7. Sedikides C, Green JD, Saunders J, Skowronski JJ, Zengel B. Mnemic neglect: Selective amnesia of one’s faults. European Review of Social Psychology. 2016 Jan 1;27(1): 1–62.

8. Swann Jr WB, Schroeder DG. The search for beauty and truth: A framework for understanding reactions to evaluations. Personality and Social Psychology Bulletin. 1995 Dec;21(12):1307–18.

9. Chang-Schneider C, Swann WB. The role of uncertainty in self-evaluative processes: Another look at the cognitive-affective crossfire. na; 2010.

10. Svoboda E, McKinnon MC, Levine B. The functional neuroanatomy of autobiographical memory: a meta-analysis. Neuropsychologia. 2006 Jan 1;44(12):2189–208.

11. Northoff G, Heinzel A, De Greck M, Bermpohl F, Dobrowolny H, Panksepp J. Self-referential processing in our brain—a meta-analysis of imaging studies on the self. Neuroimage. 2006 May 15;31(1):440–57.

12. Bird CM, Keidel JL, Ing LP, Horner AJ, Burgess N. Consolidation of complex events via reinstatement in posterior cingulate cortex. Journal of Neuroscience. 2015 Oct 28;35(43): 14426–34.

13. van der Meer L, Costafreda S, Aleman A, David AS. Self-reflection and the brain: a theoretical review and meta-analysis of neuroimaging studies with implications for schizophrenia. Neuroscience & Biobehavioral Reviews. 2010 May 1;34(6):935–46.

14. Spreng RN, Mar RA, Kim AS. The common neural basis of autobiographical memory, prospection, navigation, theory of mind, and the default mode: a quantitative metaanalysis. Journal of cognitive neuroscience. 2009 Mar;21(3):489–510.

15. Yarkoni T, Poldrack RA, Nichols TE, Van Essen DC, Wager TD. Large-scale automated synthesis of human functional neuroimaging data. Nature methods. 2011 Aug;8(8):665.

16. Amodio DM, Frith CD. Meeting of minds: the medial frontal cortex and social cognition. Nature Reviews Neuroscience. 2006 Apr;7(4):268.

17. Hughes BL, Beer JS. Protecting the self: The effect of social-evaluative threat on neural representations of self. Journal of Cognitive Neuroscience. 2013 Apr;25(4):613–22.

18. Rameson LT, Satpute AB, Lieberman MD. The neural correlates of implicit and explicit self-relevant processing. NeuroImage. 2010 Apr 1;50(2):701–8.

19. Ochsner KN, Ray RR, Hughes B, McRae K, Cooper JC, Weber J, Gabrieli JD, Gross JJ. Bottom-up and top-down processes in emotion generation: common and distinct neural mechanisms. Psychological science. 2009 Nov;20(11):1322–31.

20. Eisenberger NI, Inagaki TK, Muscatell KA, Byrne Haltom KE, Leary MR. The neural sociometer: brain mechanisms underlying state self-esteem. Journal of cognitive neuroscience. 2011 Nov;23(11):3448–55.

21. Somerville LH, Kelley WM, Heatherton TF. Self-esteem modulates medial prefrontal cortical responses to evaluative social feedback. Cerebral Cortex. 2010 Mar 29;20(12):3005–13.

22. Buckner RL, Carroll DC. Self-projection and the brain. Trends in cognitive sciences. 2007 Feb 1;11(2):49–57.

23. Leitner JB, Hehman E, Jones JM, Forbes CE. Self-enhancement influences medial frontal cortex alpha power to social rejection feedback. Journal of cognitive neuroscience. 2014 Oct;26(10):2330–41.

24. Minear M, Park DC. A lifespan database of adult facial stimuli. Behavior Research Methods, Instruments, & Computers. 2004 Nov 1;36(4):630–3.

25. Ricanek K, Tesafaye T. Morph: A longitudinal image database of normal adult age-progression. In Automatic Face and Gesture Recognition, 2006. FGR 2006. 7th International Conference on 2006 Apr 2 (pp. 341–345). IEEE.

26. Hehman E, Leitner JB, Deegan MP, Gaertner SL. Facial structure is indicative of explicit support for prejudicial beliefs. Psychological science. 2013 Mar;24(3):289–96.

27. Wickens CD. Multiple resources and performance prediction. Theoretical issues in ergonomics science. 2002 Jan 1;3(2):159–77.

28. Rosenberg M. Rosenberg self-esteem scale (RSE). Acceptance and commitment therapy. Measures package. 1965;61:52.

29. Somerville LH, Heatherton TF, Kelley WM. Anterior cingulate cortex responds differentially to expectancy violation and social rejection. Nature neuroscience. 2006 Aug;9(8):1007.

30. Scherg M. Fundamentals of dipole source potential analysis. Auditory evoked magnetic fields and electric potentials. Advances in audiology. 1990;6:40–69.

31. Scherg M. Functional imaging and localization of electromagnetic brain activity. Brain topography. 1992 Dec 1;5(2): 103–11.

32. Grech R, Cassar T, Muscat J, Camilleri KP, Fabri SG, Zervakis M, Xanthopoulos P, Sakkalis V, Vanrumste B. Review on solving the inverse problem in EEG source analysis. Journal of neuroengineering and rehabilitation. 2008 Dec;5(1):25.

33. Jentzsch I, Sommer W. Sequence-sensitive subcomponents of P300: Topographical analyses and dipole source localization. Psychophysiology. 2001 Jul;38(4):607–21.

34. Tang H, Crain S, Johnson BW. Dual temporal encoding mechanisms in human auditory cortex: Evidence from MEG and EEG. NeuroImage. 2016 Mar 1;128:32–43.

35. Santiago-Rodríguez E, Harmony T, Fernández-Bouzas A, Hernández A, Martínez-López M, Graef A, García JC, Silva-Pereyra J, Fernández T. EEG source localization of interictal epileptiform activity in patients with partial complex epilepsy: comparison between dipole modeling and brain distributed source models. Clinical Electroencephalography. 2002 Jan;33(1):42–7.

36. Scherg M, Berg P. Use of prior knowledge in brain electromagnetic source analysis. Brain topography. 1991 Dec 1;4(2):143–50.

37. Hanslmayr S, Pastötter B, Bäuml KH, Gruber S, Wimber M, Klimesch W. The electrophysiological dynamics of interference during the Stroop task. Journal of Cognitive Neuroscience. 2008 Feb;20(2):215–25.

38. Behrens TE, Hunt LT, Rushworth MF. The computation of social behavior. science. 2009 May 29;324(5931): 1160–4.

39. Saxe R. Uniquely human social cognition. Current opinion in neurobiology. 2006 Apr 1;16(2):235–9.

40. Molenberghs P, Johnson H, Henry JD, Mattingley JB. Understanding the minds of others: A neuroimaging meta-analysis. Neuroscience & Biobehavioral Reviews. 2016 Jun 1;65:276–91.

41. Allison T, Puce A, McCarthy G. Social perception from visual cues: role of the STS region. Trends in cognitive sciences. 2000 Jul 1;4(7):267–78.

42. Adolphs R. The neurobiology of social cognition. Current opinion in neurobiology. 2001 Apr 1;11(2):231–9.

43. Eisenberger NI, Lieberman MD. Why rejection hurts: a common neural alarm system for physical and social pain. Trends in cognitive sciences. 2004 Jul 31;8(7):294–300.

44. Van Overwalle F. Social cognition and the brain: a metaanalysis. Human brain mapping. 2009 Mar 1;30(3):829–58.

45. Grosbras MH, Beaton S, Eickhoff SB. Brain regions involved in human movement perception: A quantitative voxel-based metaanalysis. Human brain mapping. 2012 Feb 1;33(2):431–54.

46. Iacoboni M, Lieberman MD, Knowlton BJ, Molnar-Szakacs I, Moritz M, Throop CJ, Fiske AP. Watching social interactions produces dorsomedial prefrontal and medial parietal BOLD fMRI signal increases compared to a resting baseline. Neuroimage. 2004 Mar 1;21(3):1167–73.

47. Rotge JY, Lemogne C, Hinfray S, Huguet P, Grynszpan O, Tartour E, George N, Fossati P. A meta-analysis of the anterior cingulate contribution to social pain. Social Cognitive and Affective Neuroscience. 2014 Aug 19;10(1): 19–27.

48. Maddock RJ, Garrett AS, Buonocore MH. Remembering familiar people: the posterior cingulate cortex and autobiographical memory retrieval. Neuroscience. 2001 Jun 14;104(3):667–76.

49. Mitchell JP, Banaji MR, MacRae CN. The link between social cognition and self-referential thought in the medial prefrontal cortex. Journal of cognitive neuroscience. 2005 Aug;17(8):1306–15.

50. Sehatpour P, Molholm S, Javitt DC, Foxe JJ. Spatiotemporal dynamics of human object recognition processing: an integrated high-density electrical mapping and functional imaging study of “closure” processes. Neuroimage. 2006 Jan 15;29(2):605–18.

51. Papp N, Ktonas P. Critical evaluation of complex demodulation techniques for the quantification of bioelectrical activity. Biomedical sciences instrumentation. 1977; 13:135.

52. Sauseng P, Klimesch W. What does phase information of oscillatory brain activity tell us about cognitive processes?. Neuroscience & Biobehavioral Reviews. 2008 Jul 1;32(5): 1001–13.

53. Cohen MX. Analyzing neural time series data: theory and practice. MIT press; 2014 Jan 17. Bastiaansen M, Mazaheri A, Jensen O. Beyond ERPs: oscillatory neuronal dynamics. In The Oxford handbook of event-related potential components 2012 (pp. 31–50). Oxford University Press.

54. Hoechstetter K, Bornfleth H, Weckesser D, Ille N, Berg P, Scherg M. BESA source coherence: a new method to study cortical oscillatory coupling. Brain topography. 2004 Dec 1;16(4):233–8.

55. Forbes CE, Leitner JB. Stereotype threat engenders neural attentional bias toward negative feedback to undermine performance. Biological psychology. 2014 Oct 1;102:98–107.

56. Leary MR, Tambor ES, Terdal SK, Downs DL. Self-esteem as an interpersonal monitor: The sociometer hypothesis. Journal of personality and social psychology. 1995 Mar;68(3):518.

57. Benjamini Y, Hochberg Y. Controlling the false discovery rate: a practical and powerful approach to multiple testing. Journal of the royal statistical society. Series B (Methodological). 1995 Jan 1:289–300.

58. Singh AK, Phillips S. Hierarchical control of false discovery rate for phase locking measures of EEG synchrony. NeuroImage. 2010 Mar 1;50(1):40–7.

59. Pintzinger NM, Pfabigan DM, Pfau L, Kryspin-Exner I, Lamm C. Temperament differentially influences early information processing in men and women: preliminary electrophysiological evidence of attentional biases in healthy individuals. Biological psychology. 2017 Jan 1;122:69–79.

60. Ewald A, Aristei S, Nolte G, Rahman RA. Brain oscillations and functional connectivity during overt language production. Frontiers in psychology. 2012 Jun 7;3:166.

61. Hayes AF. PROCESS: A versatile computational tool for observed variable mediation, moderation, and conditional process modeling.

62. Buzsáki G, Draguhn A. Neuronal oscillations in cortical networks. science. 2004 Jun 25;304(5679): 1926–9.

63. Klimesch W. EEG alpha and theta oscillations reflect cognitive and memory performance: a review and analysis. Brain research reviews. 1999 Apr 1;29(2-3):169–95.

64. Kopell N, Ermentrout GB, Whittington MA, Traub RD. Gamma rhythms and beta rhythms have different synchronization properties. Proceedings of the National Academy of Sciences. 2000 Feb 15;97(4):1867–72.

65. Buzsaki G. Rhythms of the Brain. Oxford University Press; 2006 Aug 3.

